# Effective presence of antibodies against common human coronavirus in IgG immunoglobulin medicinal products

**DOI:** 10.1101/2021.07.23.453571

**Authors:** José María Díez, Carolina Romero, Rodrigo Gajardo

## Abstract

**Introduction:** In this series of studies, immunoglobulin products (IgG) formulated for different routes of administration (IV, IM, SC) and prepared from geographically diverse plasma pools were tested for activity against common human coronaviruses (HCoV). IgG products from plasma obtained from Germany, Czech Republic, Slovak Republic, USA and Spain were tested for antibodies to four common HCoV: 229E, OC43, NL63 and HKU1. Since these products are manufactured from pooled plasma from thousands of donors, the antibodies therein are a representation of the HCoV exposure of the population at large.

**Methods:** IgG products of different concentrations manufactured from geographically diverse plasma pools were tested for antibodies to four common HCoV by ELISA. In addition, neutralization assays were conducted using HCoV-229E expressed in MRC5 cells. Complete concentration-neutralization curves were obtained to calculate potencies.

**Results:** The ELISA assays showed that when expressed as specific activity (anti-HCoV activity/mg IgG) similar activity against the four common HCoV was seen across the IgG products regardless of concentration or geographic origin. Highest anti-HCoV activity was seen against HCoV-229E, followed by HCoV-OC43 and then HCoV-NL63 and HCoV-HKU1. The neutralization assays showed similar potency for two preparations of IgG prepared by different processes.

**Conclusions:** These studies are the first demonstration of antibodies to common HCoV in IgG products. The level of activity was similar regardless of the geographic origin of the plasma pool. These antibodies demonstrated neutralization activity against HCoV-229E in MRC5 cells. These results may explain the cross-reactivity seen with pre-pandemic IgG products and SARS-CoV-2 and contribute to the variability in disease course in different patients.

## Introduction

Until the appearance of SARS-CoV-2 (severe acute respiratory syndrome coronavirus 2) pandemic, relatively little attention has been paid to the classical endemic human coronaviruses (1). Common human coronaviruses (HCoVs) are globally distributed (2). They are responsible for a large proportion of respiratory infections that are mild in most cases for immunocompetent individuals. To date, four main subtypes of common HCoVs have been identified: HCoV-229E (3), HCoV-NL63 (4), HCoV-OC43 (5) and HCoV-HKU1 (6). HCoV-229E and HCoV-OC43 were discovered in 1966 and 1967, respectively, whereas HCoV-NL63 and HCoV-HKU1 were identified in 2005. None of these viruses have been found to be maintained within an animal reservoir (7). In addition, there are two other coronaviruses with animal origin that infect humans causing limited outbreaks. SARS-CoV in China in 2002-2003 and MERS-CoV (Middle East respiratory syndrome) responsible for an ongoing outbreak of severe respiratory disease in the Middle East since 2012.

Due to the ubiquity of these viruses, antibodies against common HCoVs are expected to be widely distributed in the population. Nevertheless, to our knowledge few systematic epidemiological surveys at the population level have been performed (8) and not globally. There are studies looking at the proportion of infections in some specific groups of patients (9, 10). Since a large proportion of the infections are in the childhood, whether the antibodies persist in the adult population and at what magnitude are not well known. Moreover, distinct antibody reservoirs against endemic human coronaviruses in children and adults have been described (11). Because purified medicinal immunoglobulin solutions are polyvalent and are prepared from donor plasma pools from thousands of individuals, they cover a broad spectrum of immunity in the general population and would be expected to include anti-coronavirus antibodies reflecting both the proportion of infections caused by each subtype and the specific antibody titer in the donor (general) population.

It is important to note that coronaviruses in the same subgroup, particularly betacoronavirus such as HCoV-OC43, HCoV-HKU1, SARS-CoV, SARS-CoV-2 and MERS-CoV, show some interactivity in antigenic responses. Cross reactivity between SARS-CoV and MERS-CoV with other human betacoronaviruses has become apparent (12–14). The fact that the new betacoronavirus SARS-CoV-2 is directly related to SARS-CoV (they share more than 90% sequence homology) (15) suggests that antigenic interactivity between them is possible, at least for some proteins. In addition, recently, reactions to SARS-CoV-2 in pre-pandemic immunoglobulin solutions have been observed (16). Furthermore, these solutions have some neutralizing capacity (17).

In this study, immunoglobulin (IgG) solutions for intravenous, intramuscular and subcutaneous administration were analyzed for the presence of antibodies to common HCoV. This project was designed to detect, for the first time, common HCoV antibodies in immunoglobulin IgG solutions. The immunoglobulins solutions were obtained from plasma from different origins (Germany, Czech Republic, Slovak Republic, USA and Spain) allowing an indirect comparison of the epidemiology of these viruses in these geographical areas.

## Materials and Methods

### Immunoglobulin Products

The immunoglobulin solutions used in this study were all produced by Grifols (Barcelona, Spain and Research Triangle Park, NC, USA). They included intravenous solutions (Flebogamma® DIF 5% and 10% and Gamunex®-C 10%), intramuscular solutions (Gamastan® 15-18% and Igamplia® 16%) and a subcutaneous solution (Xembify® 20%). These products were obtained from plasma pools from different origin (Germany, Czech Republic, Slovak Republic, USA and Spain).

### Immunoassays for IgG

Antibodies (IgG) to the common coronaviruses were detected using ELISA kits (Alpha Diagnostic Intl., San Antonio, TX, USA). For the α-coronaviruses the following kits were used: RV-406100 Recombivirus Human anti-HCoV 229E S1 IgG ELISA Kit and RV-406115 Recombivirus Human anti-HCoV NL63 S1 IgG ELISA Kit, For the β-coronaviruses, these kits were used: RV-406130 Recombivirus Human anti-HCoV OC43 Spike IgG ELISA Kit and RV-406145 Recombivirus Human anti-HCoV HKU1 S1 IgG ELISA Kit. The ELISAs were performed according to the manufacturer’s instructions. Data were analyzed as suggested by the kit manufacturer. The antibody potency was calculated multiplying positivity ratio for the inverse of the most diluted sample.

### Neutralization Assays

Neutralization assays was performed using HCoV-229E coronavirus. Briefly, different immunoglobulin solutions (Flebogamma ® DIF and Gamunex®-C) were incubated with 100 infectious units of the 229E virus for 1.5 hours at 37 ± 2 °C. MRC5 cells (ATCC CCL-171™, Manassas, VA, USA) in confluent culture in 96-well microtiter plates were infected with 200 μL per well of virus/antibody mixture. The microtiter plates were incubated at 35 ± 2 °C for 4 days and cytopathic effects were observed using an inverted microscope (Axiovert 40, ACHROPLAN 10X/0.25 Ph1 objective, Karl Zeiss, Göttingen, Germany). Concentration-effect curves were generated and IC_50_ values were calculated using a GraphPad Prism software (Version 9.1.0 for Windows, GraphPad Software, San Diego, California USA).

## Results

The IgG titers (anti-coronavirus activity/mL) for the immunoglobulin products are shown in Figure 1. When expressed in this manner, the lower concentration of immunoglobulins (5%) showed less activity than the higher concentrations (10-20%). For products of similar concentration, IgG activity was similar regardless of the geographic origin of the plasma pool. Overall, the highest activity was seen against the HCoV-229E and HCoV-OC43 strains.

**Figure 1:**
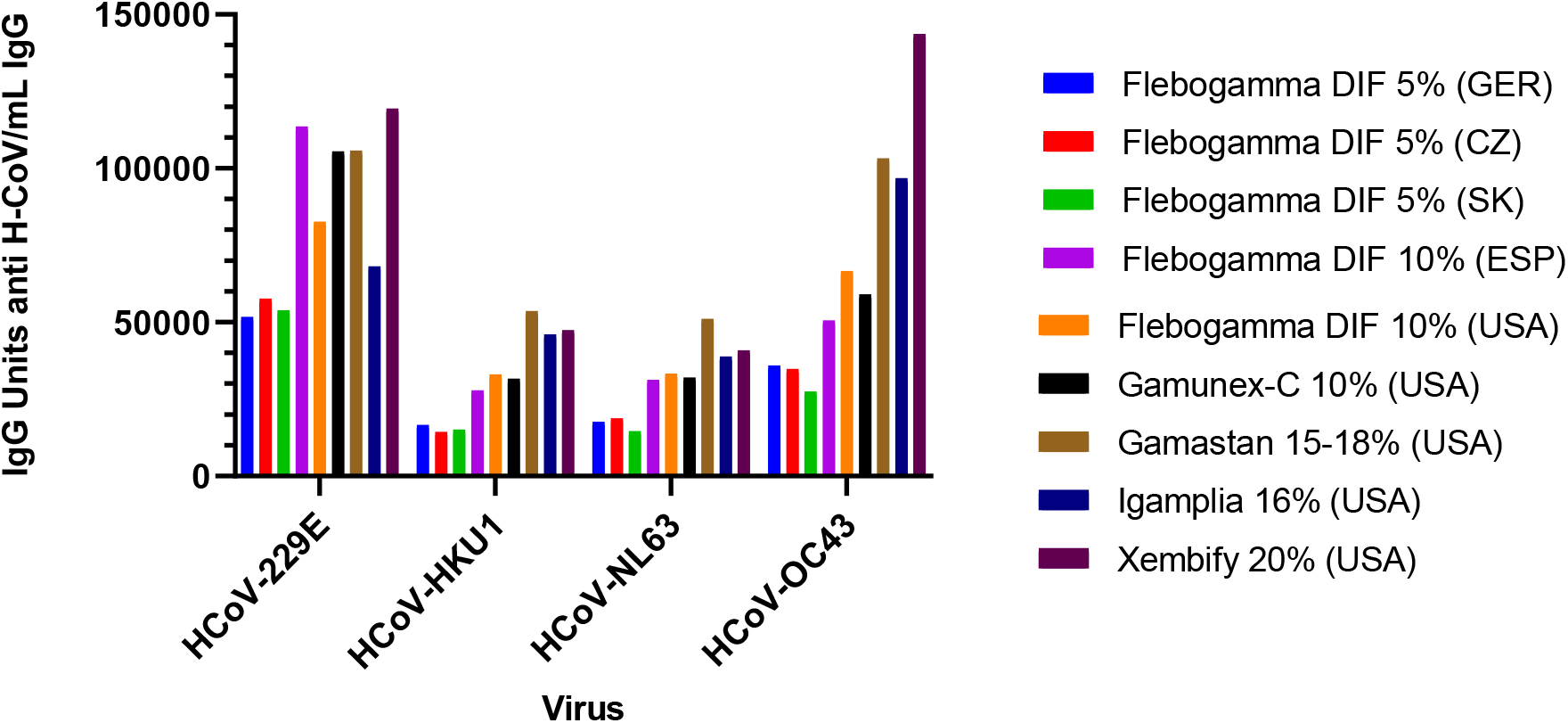
IgG against human common coronaviruses (HCoV) per product mL. Anti-HCoV activity (as measured by IgG ELISA in units of anti-coronavirus IgG/mL of product) to common coronaviruses in different immunoglobulin solutions manufactured using plasma from different countries. The levels of antibodies (IgG) against the same virus were similar in all products with a similar IgG concentration. However, differences were seen between the viruses.

The similarity becomes clearer when the data were expressed as specific activity (anti-coronavirus activity/mg IgG: Figure 2). These data show that anti-coronavirus activity was consistent across the products regardless of the total IgG concentration and the origin of the plasma pool. Activity was highest against the HCoV-229E followed by HCoV-OC43. Similar lower level activity was seen against HCoV-NL63, and HCoV-HKU1.

**Figure 2:**
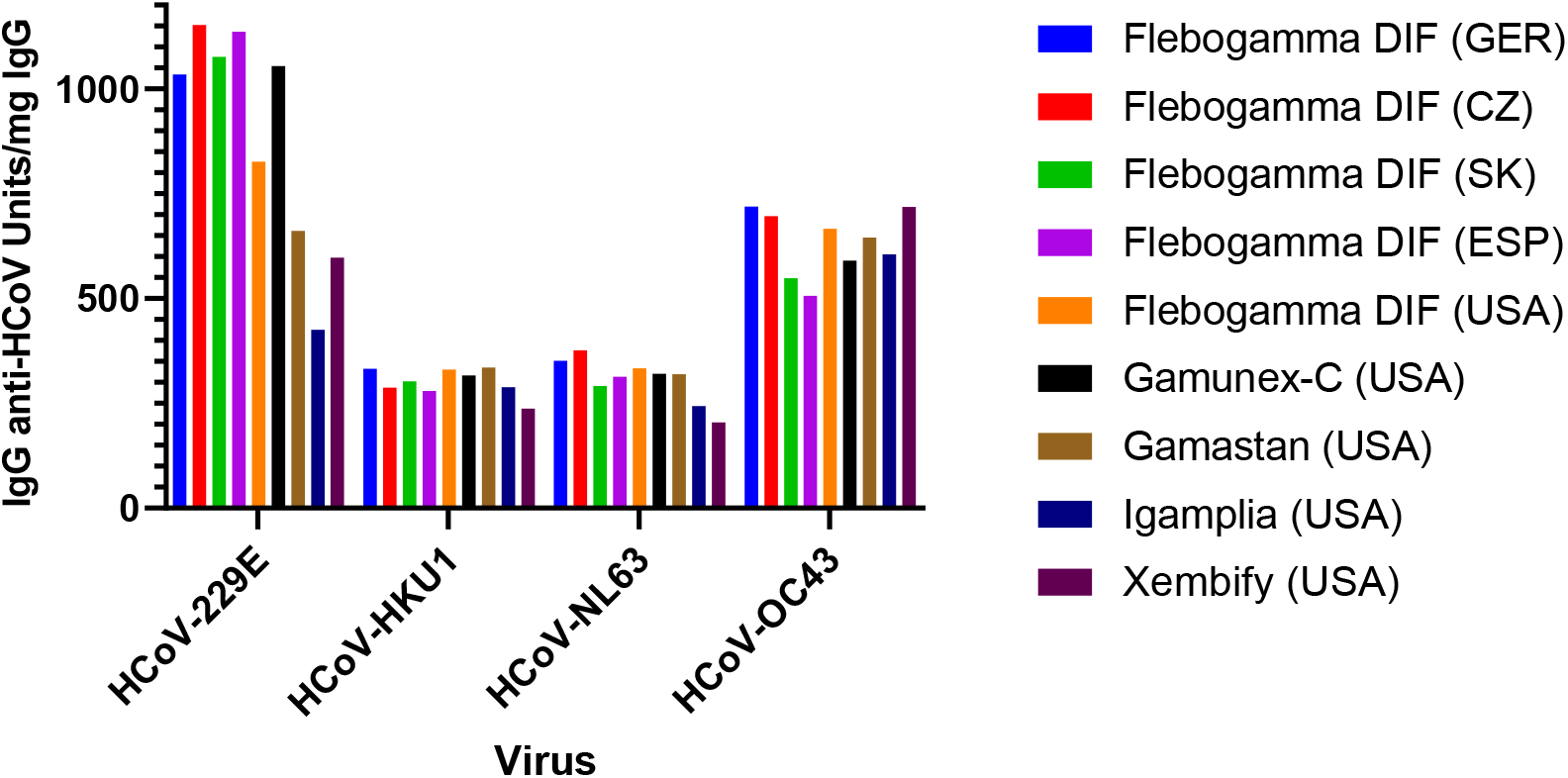
IgG against human common coronaviruses (HCoV) per mg whole IgG Anti-HCoV activity measured by IgG ELISA (expressed as units of anti-HCoV IgG/mg total IgG) against common HCoVs. Specific activity of the anti-HCoV antibodies was similar regardless of the geographic origin of the plasma pool.

When the data from all the products was combined, the mean specific activity against the individual virus strains (Figure 3) followed the same profile as that noted for the individual products (Figure 2). Greatest activity was seen against the HCoV-229E (885 ± 267units anti-HCoV activity/mg IgG) virus followed by the HCoV-OC43 virus (633 ± 76 units anti-HCoV activity/mg IgG) with similar lower levels of activity observed against the HCoV-NL63 (306 ± 53 units anti-HCoV activity/mg IgG) and HCoV-HKU1viruses (301 ± 32 units anti-HCoV activity/mg IgG).

**Figure 3:**
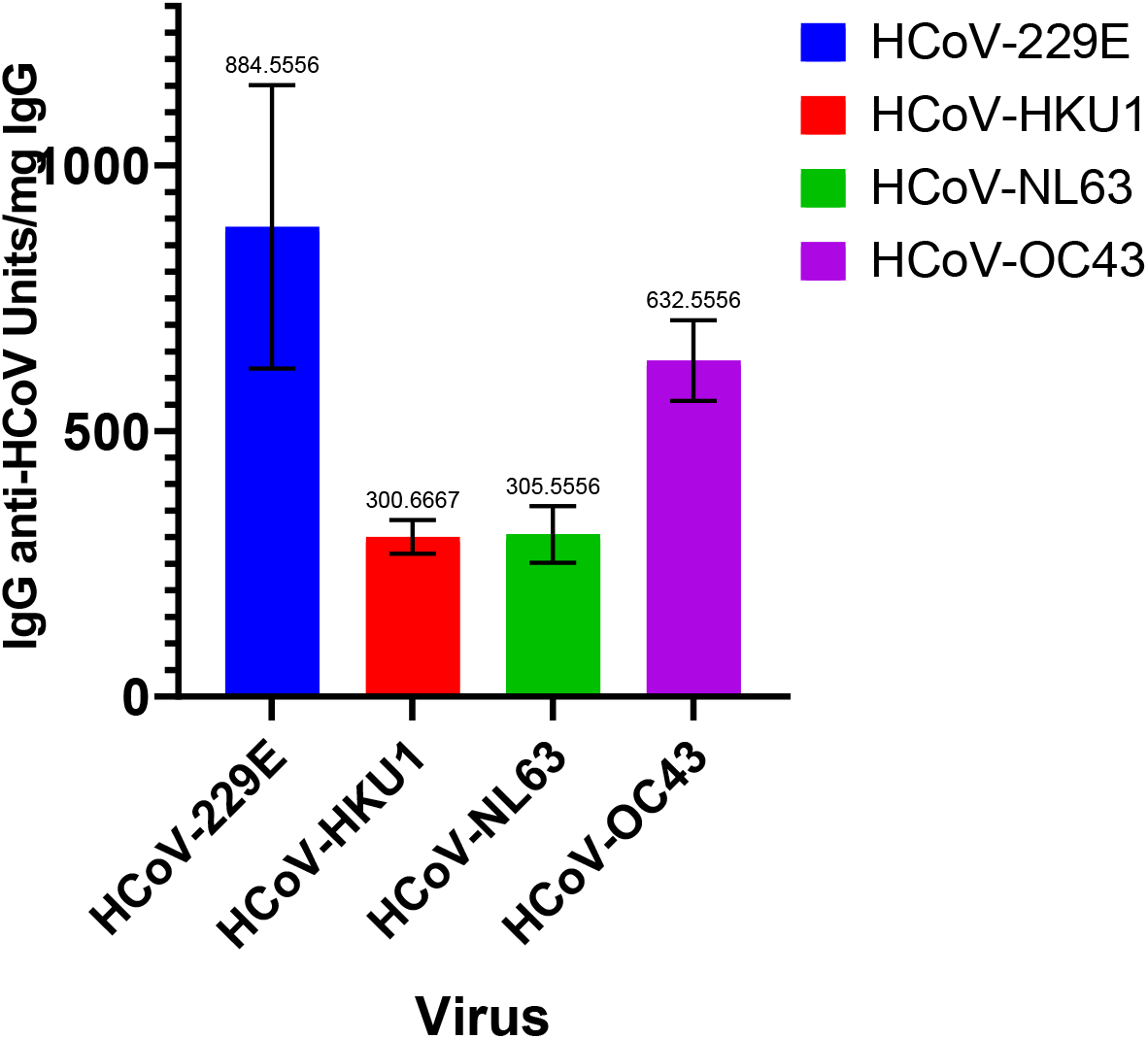
Levels IgG against common human coronaviruses (HCoV) in product average. Mean anti-HCoV antibody levels (measured by IgG ELISA and expressed as units/ mg IgG) across all products were different for each virus (ANOVA, p < 0.0001) except for HCoV-HKU1 and HCoV-NL63 which show similar antibody levels.

IgG activity results were also analyzed after segregating the results by geographic plasma origin into three groups: central Europe (Czech Republic and Slovak Republic), Spain and USA (Figure 4). IgG products had similar activity against all four HCoV regardless of the geographic origin of the plasma.

**Figure 4:**
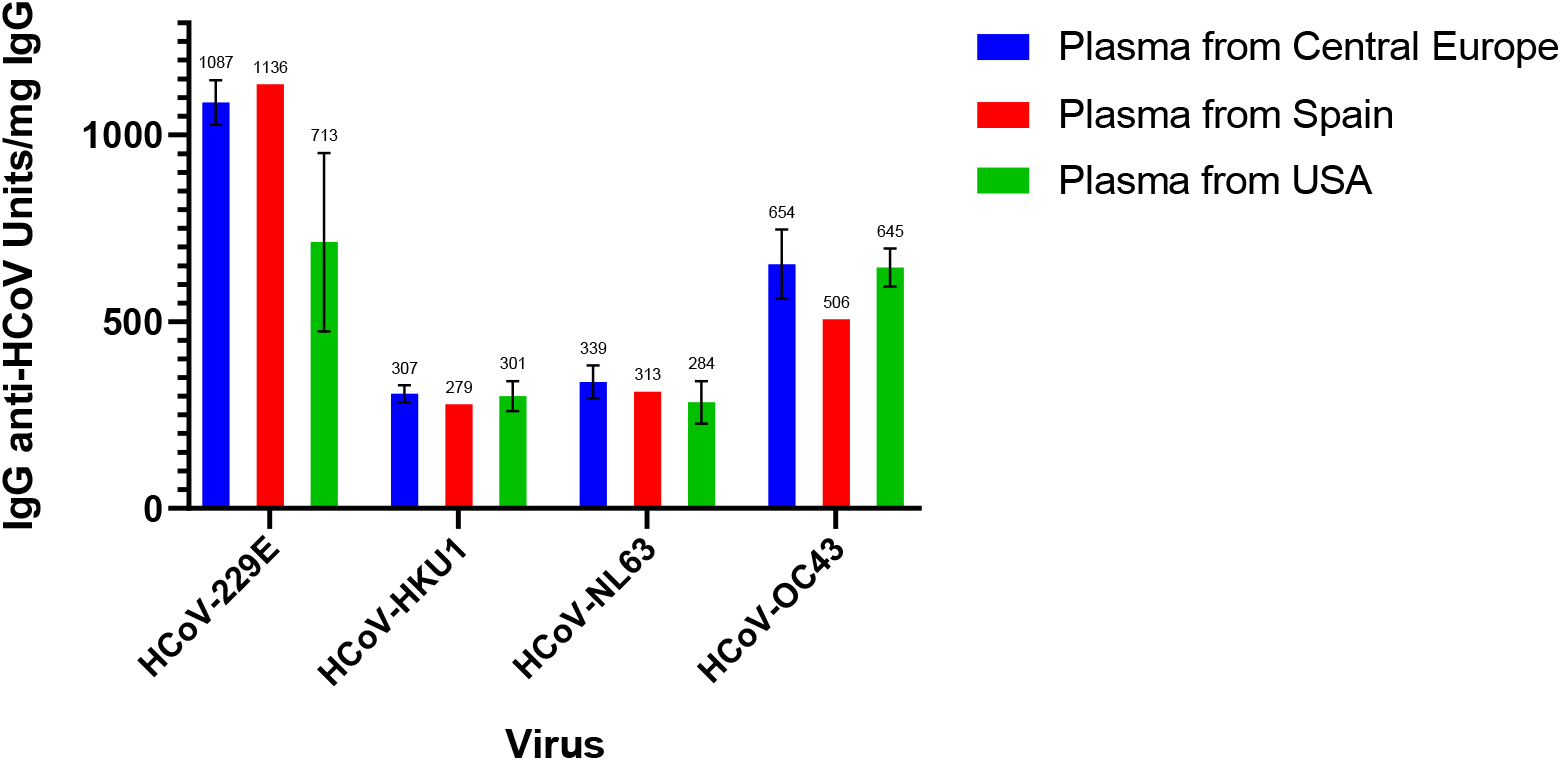
Antibodies to common human coronaviruses (HCoV) by plasma origin. Antibody levels (measured by IgG ELISA and expressed as units of anti-HCoV activity/mg IgG) to human common coronaviruses grouped by geographic origin of the plasma pool. Differences were seen among the common HCoV strains studied, but there were no statistically significant differences between product derived from plasma of different geographic origins (p value 0.8951).

**Figure 5:**
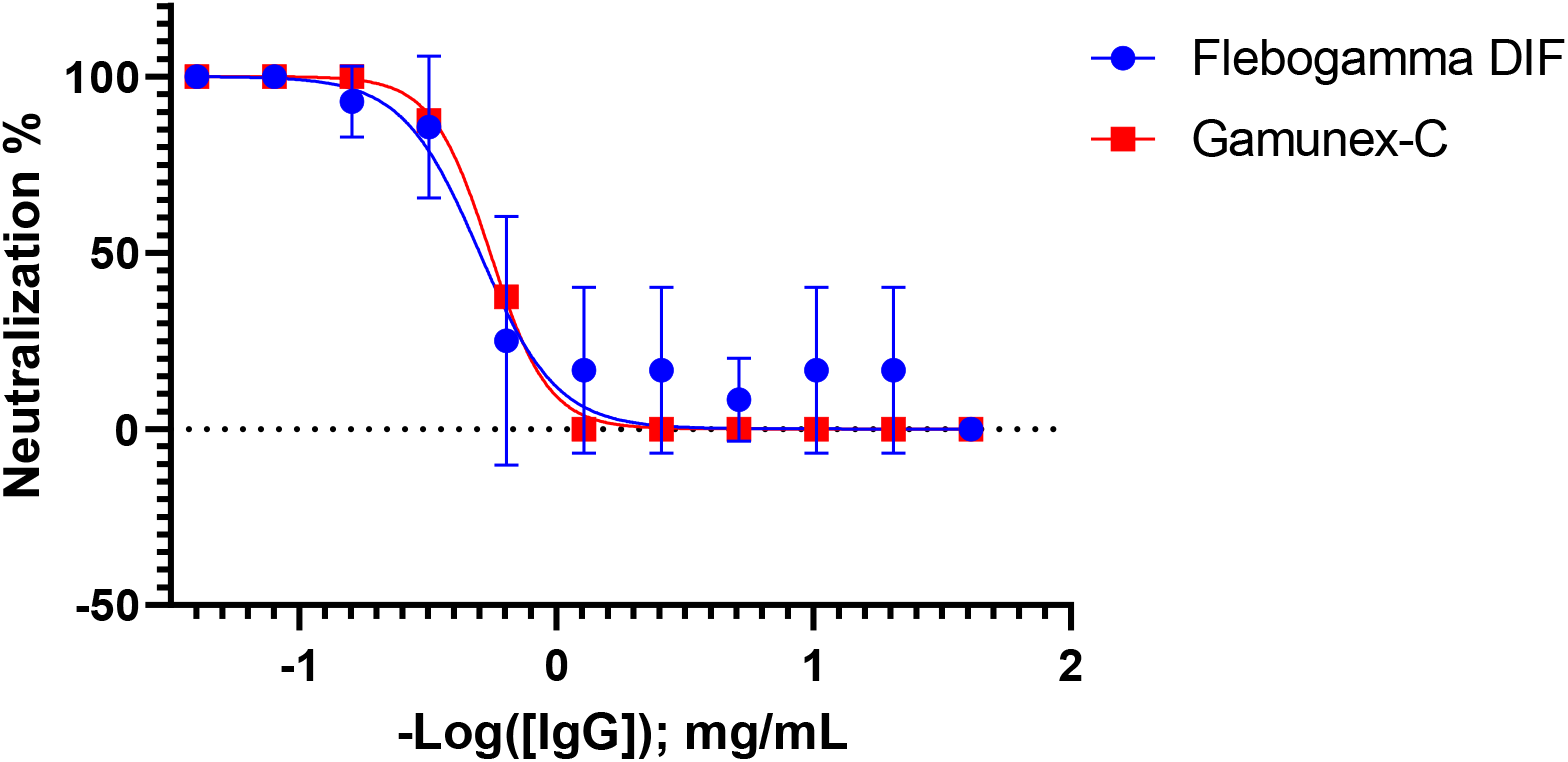
HCoV-229E virus neutralization. Neutralization of HCoV-229E was measured in a cytopathic assay in MRC5 cells. Concentration-effect curves (mg IgG/mL-neutralization %) were generated for virus neutralization and IC_50_ values were calculated. The IC_50_ for Flebogamma®-DIF was 0.503 mg IgG/mL which was very similar to the IC_50_ for Gamunex®-C (0.553 mg IgG/mL).

Functional characterization of these antibodies was performed by infectivity neutralization assays using HCoV-229E. When neutralization assays were performed using the HCoV-229E virus in MRC5 cells, the concentration-effect curves for Flebogamma®-DIF (10 %, origin: USA) and Gamunex®-C (10%, origin: USA) were nearly superimposable. This shows that the neutralization activity of the antibodies present in these products is essentially the same regardless the manufacturing process. This demonstrates that the IgG medicinal products contain functional antibodies against common HCoVs.

## Discussion

For the first time, the presence of antibodies to common coronaviruses was measured in therapeutic immunoglobulin solutions (IVIG, IMIG, and SCIG). Anti-HCoV IgG levels were similar across products for each virus regardless of the product concentration or the geographic origin of the plasma. However, there are differences in antibody levels between viruses with higher levels of IgG HCoV-229E, with lower levels for HCoV-OC43, and similar still lower levels for HCoV-HKU1 and HCoV-NL63. Studies on the incidence of HCoV have shown that the most common strain and prevalence depend on the geographic region and the time of year.

Gaunt et al. found that the most prevalent strain of common HCoV in Edinburgh, Scotland varied from year to year and that respiratory infections due to common HCoVs showed marked seasonality. However, over the three-year data collection period, HCoV-OC43 and HCoV-NL63 were the most frequently detected common HCoVs. (9) Similar seasonality and variation in the predominant viral strain from year to year was also found in a study conducted in the United States (8).

A study in France found that HCoV-229E and HCoV-HKU1 were the most common HCoVs causing respiratory infections. (18) In Japan, HCoV infections were most commonly caused by HCoV-NL63 and HCoV-HKU1 with peak prevalence in the winter months and annual variation in the relative prevalence of the different common HCoV strains. (19) One pediatric study in China found that HCoV-229E and HCoV-OC43 had the highest prevalence of the common strains for causing respiratory infections (20) while another study found HCoV-NL63 to be the most prevalent. (21) Co-infection with other respiratory viruses was also a common finding (9, 18, 20).

A global, systematic review and meta-analysis of data from 1995-2020 in pediatric and adult patients showed that OC43 was the most prevalent common HCoV (estimated prevalence 2.40%) followed by NL63 (1.60%), HKU1 (1.04%) and 229E (0.97%). These data were collected almost exclusively in developed countries (97%) (1).

Given the above studies showing differences in the prevalence of common HCoV strains in different parts of the world, it is somewhat surprising that all the IG samples in this study showed a similar pattern of anti-HCoV activity. This could be explained by the seasonal variability of the prevalence of the common HCoVs, i.e., the predominance of one strain in a given winter season followed by the predominance of a different strain in the following winter season, and that the plasma pool likely reflects HCoV exposure over time in the donors. In addition, two of the epidemiological studies previously cited were conducted in Asia while the IG products tested in this study were from Eastern Europe, Spain and the USA. The predominance of different HCoV strains varies in different geographical areas over time.

It is also unexpected that the antibody profile in the IG products does not match the HCoV prevalence in the longitudinal meta-analysis. This may reflect that the geographic source of the plasma used to produce these products is reflective of these specific regions and not representative of worldwide prevalence. Another factor that could contribute to the apparent disparity may be that the published studies represent clinical samples from patients that sought medical attention while the IG products represent a population that included individuals who had milder infections and did not seek medical attention. In other words, the epidemiology reflects patients with more symptomatic infections while the IG products include individuals that had asymptomatic or mild infections as well as symptomatic infections.

In addition, these studies demonstrated that these antibodies had neutralizing activity against HCoV-229E virus in MRC5 cells. Neutralization activity is crucial to the use of plasma-derived product used in the treatment and/or prevention of viral diseases. The neutralizing capacity in this study was demonstrated with two different products with different manufacturing methods. This finding suggests the ubiquity of anti-HCoV binding activity is accompanied by neutralization activity.

It is also important to note that coronaviruses in the same subgroup, particularly betacoronaviruses such as HCoV-OC43, HCoV-HKU1, SARS-CoV, SARS-CoV-2 and MERS-CoV, show some interactivity in antigenicity. Cross-reactivity between SARS-CoV and MERS-CoV and other human betacoronaviruses has been reported (12–14). The fact that the recently identified betacoronavirus, SARS-CoV-2, is closely related to SARS-CoV (> 90% sequence homology) (15) suggests that antigenic interactivity between them is possible, at least for some proteins.

In addition, reactivity to SARS-CoV-2 in pre-pandemic immunoglobulin solutions has recently been observed (16). As demonstrated in this study, these solutions also have the capacity to neutralize common HCoVs such as HCoV-229E. Furthermore, these solutions have demonstrated some neutralizing capacity towards SARS-CoV-2 (17). The worldwide presence of these common HCoVs may affect the current SARS-CoV-2 pandemic. Pre-existing immunity to common HCoVs may have a role in both humoral and cellular responses to SARS-CoV-2 (17, 22, 23), and it could explain, in part, the differences in illness behavior among patients.

In conclusion, these studies demonstrated for the first time the presence of antibodies to common HCoVs in parenteral IG products. The level of anti-HCoV activity for each virus was similar regardless of the geographic origin of the plasma pool. Neutralization activity was demonstrated against a representative strain of HCoV (HCoV-229E) in MRC5 cells. These findings may help to explain the previously evidenced cross-reactivity and neutralization activity for SARS-CoV-2 observed with pre-pandemic IG products (16, 17), and differences in illness behavior among patients.

## Acknowledgments

Michael K James Ph.D. (Grifols) is acknowledged for medical writing and editorial support in the preparation of this manuscript. The authors acknowledge the expert technical assistance from D Casals, E Sala, and J Luque.

